# Unexpected variability in laboratory and clinical assays used to quantify *Cryptococcus neoformans* polysaccharide

**DOI:** 10.1101/2025.05.13.653753

**Authors:** Maggie P. Wear, Davin Kim, Risha Roy, Arturo Casadevall

## Abstract

The measurement of cryptococcal capsular polysaccharide (PS) amount and concentration is essential for laboratory studies of its structure and function. Clinically, the detection of cryptococcal capsular polysaccharide as cryptococcal antigen is a foundational assay for the diagnosis of cryptococcosis. In this study we compared the results of four laboratory assays for measuring cryptococcal polysaccharide (gravimetric determination, phenol-sulfuric acid reactivity, Size Exclusion Chromatography Multi-Angle Light Scattering (SEC-MALS), and ELISA) and two clinical assays (lateral flow assay (LFA) (IMMY) and the cryptococcal latex agglutination assay (CLAA) (Remel)) using four cryptococcal strains that each express single repeating polysaccharide motifs. The results show that the sensitivity of laboratory assays for PS was relatively consistent but manifested significant variation with strain. Phenol-sulfuric acid determination was highly variable unless the PS was pre-digested with HCl, suggesting that differences in the structural complexity of the polysaccharide could affect timing and intensity of the colorimetric reaction. Serological assays were the most sensitive, but these showed the most strain-to-strain variation on the threshold of PS detection consistent with antigenic differences. Differences in the reactivity of LFA and CLAA for cryptococcal PS from different strains imply that comparable cryptococcal antigen titers could reflect different concentrations of PS in bodily fluids. In summary, we find that variation in PS structure amongst strains can affect the sensitivity of the various tests used to measure cryptococcal PS. (Word count: 224)

## Introduction

One of the most important virulence factors of the human pathogenic fungi *Cryptococcus neoformans* and *Cryptococcus gattii* are their polysaccharides (PS), both exo-(EPS) and capsular (CPS). These PS impart a variety of defensive capabilities including enhanced resistance to antifungal compounds (1), interference with immune responses (2) and evasion of phagocytosis by macrophages (3), thus contributing to cryptococcal pathogenesis. Consisting primarily of glucuronoxylomannan (GXM), and to a lesser extent, glucuronoxylomannogalactan (GXMgal), cryptococcal PS have been identified for use in future vaccines and as targets for potential monoclonal antibody therapies (4). However, the functionality and molecular structure of these PS have not been fully elucidated. A critical step in both structural characterization and diagnosis of cryptococcal infection is accurate identification and quantification of cryptococcal PS.

There are a few methodological perspectives by which PS are quantified: physical, chemical, and ligand binding. Isolation of cryptococcal EPS, usually by cetyltrimethylammonium bromide (CTAB) precipitation which favors GXM, often involves PS quantification to determine the yield. Cryptococcal PS can be quantified gravimetrically via lyophilization to determine the dry weight of the sample after EPS precipitation (Figure 1). The fact that the entire sample is included in the dry weight has led some to question whether other biomolecules like lipids (5, 6) and proteins (2, 6) make up a significant portion of the sample. Various studies have used light scattering to measure the particle size of PS (2, 7), but when multi-angle light scattering (MALS) is coupled to size exclusion chromatography (SEC) it can measure mass along with molecular weight and shape (8)(9). SEC-MALS as a PS quantification method is further advantaged by the small quantity of sample needed, along with its ability to resolve and purify heterogeneous samples. Direct quantification of PS chemically by the phenol-sulfuric acid (PSA) assay utilizes sulfuric acid to break polymers into monomers and further into furfurals, which with phenol produce a colored reagent that can be quantified calorimetrically using a standard curve (10). While the PSA is the favored method for bacterial and plant polysaccharides (11), these PS are linear compared to the branched structure of fungal PS. Finally, serological assays utilize antibody binding to cryptococcal PS to both identify and quantify the amount of PS. Historically, enzyme-linked immunosorbent assays (ELISA) utilizing monoclonal antibodies reactive against cryptococcal PS (12) have been the primary biochemical method of PS quantification. The binding level of antibodies specific to cryptococcal PS epitopes, quantified by chemiluminescent detection allows for the construction of a standard curve from which sample concentrations are extrapolated. While highly sensitivity and specific, particularly the competitive ELISA or sandwich ELISA, this method is also highly variable due to the indirect nature of quantification (13). In addition to ELISA the two major clinical tests for cryptococcal infection, cryptococcal antigen latex agglutination (CLAA) and the lateral flow assay (LFA) rely on antibody binding to PS, the cryptococcal antigen (CrAg) for diagnosis. In this work we compare the replicability, specificity, and sensitivity of these methods for PS that differs in GXM structural motif composition. The results show considerable inter-method variability including clinical assays, which can carry important implications for laboratory studies and the interpretation of cryptococcal antigen determinations.

**Figure 1:**
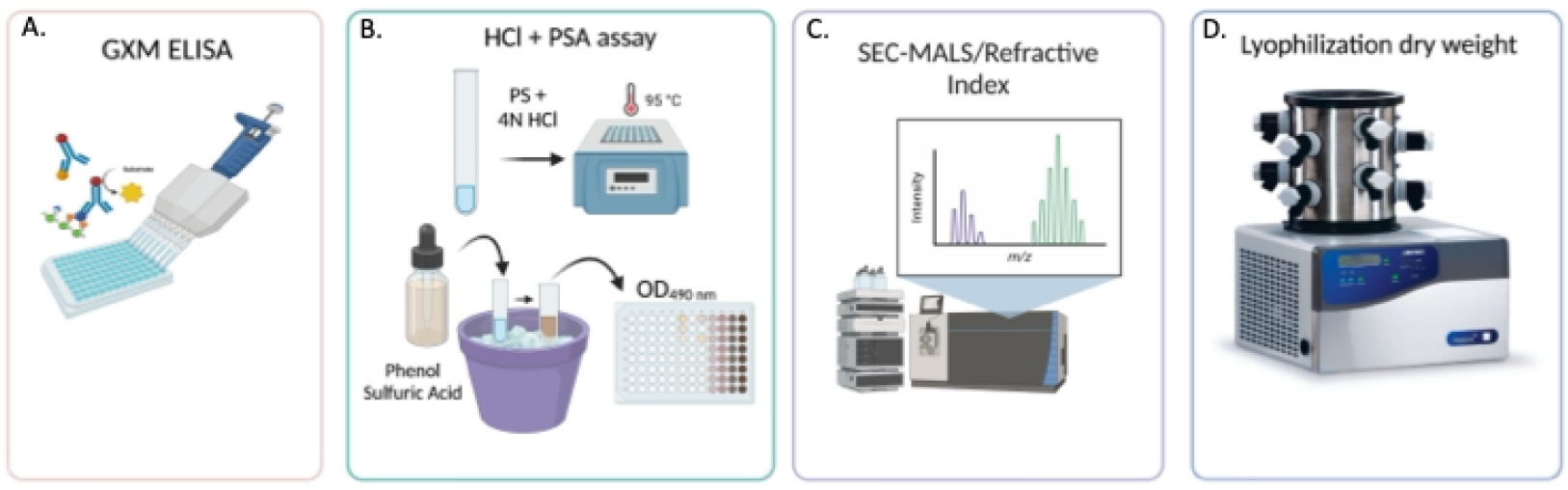
Methods used to quantify cryptococcal PS **A**. GXM ELISA uses antibodies to detect PS and quantify relative to standard. **B**. Modified PSA assay includes a pre-incubation step in 4N hydrochloric acid to reduce the complexity of cryptococcal polysaccharides prior to digestion with phenol-sulfuric acid solution. **C**. SEC-MALS uses refractive index to determine the number of populations of polysaccharide along with the molecular weight, shape (nm), and mass of each population. **D**. Dry weight by lyophilization is a gravimetric determination of the amount of polysaccharide present in a sample.

## Methods

### Cryptococcal Strains and PS

Each quantification method was compared using exopolysaccharide (EPS) and capsular polysaccharide (CPS) samples from single motif strains expressing solely the M1, M2, M3, or M4 GXM structural motifs (14), corresponding to the 24067, mu-1, 409, and kt24066 strains, respectively. EPS samples were collected as previously described (15). Following filtration of the supernatant through a 0.22 µm filter, the resultant PS samples were not fractionated and were kept whole; samples collected in this way are referred to as ‘whole EPS’ or wEPS. To remove media components, salt, and other small molecules wEPS was filtered through a 3 kDa molecular weight cut-off (MWCO) Amicon filter (Millipore Sigma, St. Louis, MO, USA). Afterwards the sample was washed with water and the >3 kDa retentate collected and referred to as EPS. CPS samples were isolated as done previously (16).

### Dry Weight

A pre-determined volume of each EPS sample was pipetted into weighted Eppendorf tubes (after filtration through a 3 kDa MWCO Amicon to remove salts and small molecules) and placed at -20°C until samples were frozen. The frozen samples were then lyophilized until dehydrated, a time determined when the measured mass did not change over 24 hours, and dry weight was determined after accounting for the empty tube mass as previously reported (17). Sample concentrations were determined by dividing the obtained dry weight by the original sample volume.

### Protein Quantification by Bradford Assay

A 5 µg/mL bovine serum albumin (BSA) solution was prepared as a protein standard from the Pierce (Waltham, MA USA) Bovine Serum Albumin Standard Pre-Diluted Set. A 160 µL volume of each of the cryptococcal PS samples at unknown protein concentration along with the BSA were combined with 40 µL of Bio-Rad Protein Assay (Hercules, CA USA) dye reagent in Eppendorf tubes and vortexed thoroughly before being plated into a microtiter plate and diluted according to a 1:2 dilution series.

The final plate was incubated for 5 minutes at room temperature before the absorbances of each well were measured via microplate reader at a wavelength of 595 nm. Using a standard curve constructed from the optical densities at BSA each concentration, each of the sample proteins concentrations were determined from the well optical densities (Supp. Fig. 1).

### SEC-MALS

SEC-MALS was run on an Agilent (Santa Clara, CA USA) 1200 HPLC supplied with a degassing module, auto-injector, a 50 µL injection loop, a diode array detector (DAD), and an in-line Wyatt (Santa Barbara, CA, USA) mini-DAWN MALS detector prior to a refractive index (RI) detector. Samples of a 25 µL injection volume each were run on an Acclaim SEC-300 analytical column (300 Å, 4.6 × 300 mm) at 0.3 mL/min flow rate of a water mobile phase for a static run of 40 minutes. The resulting data was subsequently analyzed using ASTRA, a light scattering analysis tool software offered by Wyatt. UV, light scattering (LS), and RI baselines were first normalized before peaks were selected based on correlation of these measures. Experimental sample weight average molar mass and sizes were determined by comparison to p5 and p50 standards (pullulan samples of sizes 5 kDa and 50kDa at 0.94 mg/mL and 7.57 mg/mL respectively). Dividing mass by the injection volume yielded a concentration measure for each sample run.

### PSA Assay with Modifications

Hydrolyzed samples were first prepared as described by Gao and colleagues (18). However, given our findings from the time course implying that GXM structural complexity required longer reaction times, this was subsequently altered depending on sample identity, as single-motif EPS samples were incubated for 30 minutes while single-motif CPS samples were incubated for 50 minutes. Following HCl pre-treatment and incubation, samples were incubated 95°C for 30 minutes in freshly prepared phenol:water:sulfuric acid (0.1:0.5:1.0, w:v:v) before cooling on for 5 minutes. A D-mannose monosaccharide standard (10 mg/mL) was included and sample and standard absorbances at 490 nm were determined. Polysaccharide concentration was calculated relative to a mannose standard curve. To determine the time of HCl pretreatment necessary for accurate polysaccharide quantification PS samples were incubated for 140 minutes, with samples taken at 10-minute intervals starting at 30 minutes. Two standards were included, 30-minute HCl-pretreated H99 EPS and D-mannose standard. Following incubation, the PSA assay was performed and concentrations determined as detailed above.

### GXM ELISA

High-binding polystyrene microplates (Corning, NY USA)were first coated with goat anti-mouse IgM at 1 µg/mL in PBS (Southern Biotech Birmingham, AL USA) and then blocked in 1% BSA blocking solution prior to incubation and washing (13). Note all subsequent references to incubation denote an hour-long incubation period at 37°C, and all instances of references to plate washing refer to 3 cycles of rinses with 0.1% Tween 20 in Tris-buffered saline. Next, a murine anti-GXM IgM at 2 µg/mL in blocking buffer was added as a capture antibody, then incubated and washed. The capture antibody used varied based on the dominant motif of the GXM antigen as found using glycan arrays (19),(20): mAb 2D10 was used for GXM samples predominantly bearing the M1, M2, or M3 motifs, and mAb 12A1 was used for samples predominately bearing the M4 motif. Then, either EPS or CPS samples were added in conjunction with a H99 EPS standard of known concentration and serially diluted with blocking buffer before being incubated and washed. mAb 18B7, an unlabeled murine anti-GXM IgG1 (Southern Biotech Birmingham, AL USA), was subsequently added at 5 µg/mL in blocking buffer as a detection antibody. Following another 1-hour incubation and wash, a secondary goat anti-mouse IgG1 conjugated to alkaline phosphatase was added at 1 µg/mL in blocking buffer. After a final incubation and wash step, the assay was developed at room temperature by the addition of 1 mg/mL p-nitrophenyl phosphate (pNPP) (Thermo Scientific Waltham, MA USA) in substrate buffer. The absorbance of each well at 405 nm was measured using a microplate reader using (ID5, Molecular Devices, San Jose, CA USA), and a standard curve was constructed by plotting the obtained optical densities against the known standard concentrations. The resulting linear regression equation was referenced to obtain EPS and CPS sample concentrations.

### Lateral Flow Assay (LFA)

The Thermo Scientific (Waltham, MA USA) IMMY CrAg Lateral Flow Assay kit was used to evaluate the sensitivity of the test to cryptococcal exopolysaccharide antigen (EPS) from four single-motif expressing strains (24067, mu-1, 409, and kt24066). The LFA utilizes two antibodies, IgG3s F12D2 and 339. MAb F12D2 was raised against CTAB isolated EPS from 100% M2 expressing strain Mu-1 (21). F12D2 was found to bind CTAB isolated EPS from single motif expressing strains Mu-1, 409, 24066, and 9759 as well as de-O-acetylated GXM (21). MAb 339 was raised against CTAB isolated EPS from strain 24065, 100% M3 expressing (22) and binds CTAB isolated EPS from strain NIH-6 (67% M2, 17% M3, 15% M6), as well as single motif expressing strains 409 (M3) and 9375 (M1). To estimate the limit of detection for each strain the directions supplied with the IMMY kit were followed wherein the strip with the lowest antigen concentration that met the defined positivity criteria represents the limit of detection. To provide a quantitative assessment of positive detection the band intensity was also analyzed using ImageJ software. A positive result was determined by imaging each lateral flow strip and determining the integrated densities of the control and sample bands. Any instance where subtraction of the integrated density value of the control band from the sample band yielded a positive value was treated as a positive result indicative of antigen detection.

### Cryptococcal Latex Agglutination Assay (CLAA)

Cryptococcal exopolysaccharide (EPS) antigen titer tests were performed using Thermo Scientific Remel (Waltham, MA USA) Cryptococcus Antigen Test Kits. The CLAA uses a single IgM mAb BA4-DF5 resulting from immunization and fusion with strain 9759 (15% M1, 58% M2, 14% M5, 13% M6) which is able to bind CTAB EPS from strains 9759, 3175 (aka M9030 – 20% M1, 30% M2, 45% M6), and 1254 (62% M1, 19% M5, 19% M6) (23, 24). Preparations of EPS samples set to a uniform concentration of 5 ug/mL from the single-motif expressing 24067, mu-1, 409, and kt24066 strains along with the mixed motif KN99a strain were used to determine the limit of detection. As cryptococcal polysaccharide preparations were made in 1x Dulbecco’s Phosphate Buffered Saline (DPBS), these samples were considered more similar to cerebral spinal fluid than serum clinical specimens and therefore were not treated with protease before immersion in a boiling water bath for 5 minutes. The test kit instructions were followed and each reaction card containing the various antigen concentrations for a given strain was allowed to incubate for 5 minutes on a clinical rotator and immediately observed for agglutination. Any test circles showing latex clumps (visualized as small discrete spots of white against the black background) were deemed positive results. The limit of detection for each strain by the latex agglutination method was determined as the lowest concentration test circle or titer that yielded agglutination.

### Statistical Analysis

Results were analyzed using a two-way ANOVA with a Bonferroni correction for multiple measures where the variables were replicate and strain. The examination of sensitivity lies in the limit of detection for the quantification methods. For the clinical assays and ELISA, the limit of detection is <1 µg/mL, but the other laboratory methods quantify yield in mg/mL. Only the samples for which there is a statistically significant difference (p < 0.05) are indicated in the figures. All others did not show a statistically significant difference.

## Results

### Lyophilized cryptococcal polysaccharide contains negligible amounts of protein

A significant concern about utilizing dry weight to quantify cryptococcal PS is that these samples may contain significant amounts of protein. To determine the protein content of EPS, we performed a Bradford assay for protein quantification of samples from the four single motif expressing strains (Supp. Fig. 1). The Bradford assay was selected over the BCA assay due to BCA reagents reacting with PS (25). This analysis revealed that proteins accounted for 0.001-0.0013% of the total mass (24067: 1.31 µg protein/mg EPS; Mu-1: 1.27 µg protein/mg EPS; 409: 1.08 µg protein/mg EPS; KT24066: 1.06 µg protein/mg EPS), making a negligible compared to the PS mass. The average dry weight (of 3 biological replicates) of EPS from 24067 was 8.22 mg/mL ± 1.37, Mu-1 was 8.11 mg/mL ± 0.62, 409 was 8.33 mg/mL ± 1.19, and KT24066 was 9.56 mg/mL ± 1.34, yielding a protein content of EPS of 0.08 – 0.1%.

### SEC-MALS PS quantification requires small sample amounts and shows low variability

Size Exclusion Chromatography Multi-Angle Light Scattering (SEC-MALS) allows the separation of materials by size while also measuring molecular weight, amount, shape, and polydispersity. Utilizing this method, we observed that EPS from all four single motif expressing strains separated into three peaks by light scattering profile but only a single peak by refractive index of the solution (Figure 2). Since refractive index detects polysaccharides (26), we quantified our samples based on this peak which eluted 9 and 12 minutes of a 20-minute run (Figure 2A). Analysis of the RI peaks shows that the *C. neoformans* strains 24067 and Mu-1 preparations contained 7.62 µg/mL ± 1.51 and 4.02 µg/mL ± 2.60 of EPS while the *C. gattii* strains 409 and kt24066 preparations contained 5.08 µg/mL ± 2.64 and 6.79 µg/mL ± 1.30, respectively (Figure 2B). The size of EPS polymers varied with 24067 and Mu-1 being 210 and 121 kDa, respectively and 409 and kt24066 being smaller at 49.5 and 33.9 kDa, respectively (Figure 2C).

**Figure 2:**
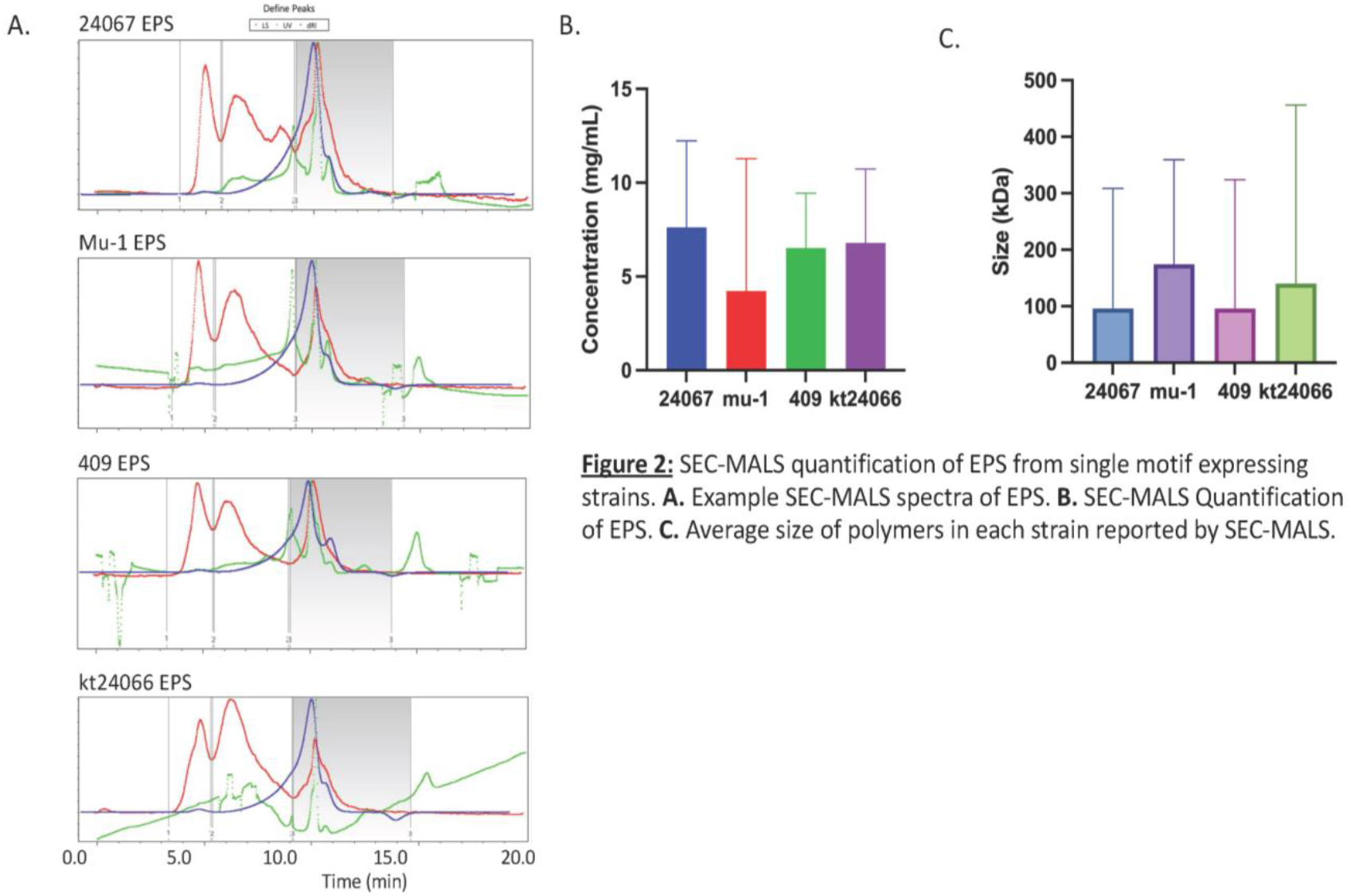
SEC-MALS quantification of EPS from single motif expressing strains. **A**. Example SEC-MALS spectra of EPS. **B**. SEC-MALS Quantification of EPS. **C**. Average size of polymers in each strain reported by SEC-MALS.

### HCl preincubation improves PSA assay quantification

The standard PSA assay resulted in a significantly lower PS concentration than dry weight or SEC-MALS measurements (data not shown). GXM macromolecules have been proposed to form dendrimers with a dense core to which fibrils are attached (27). Consequently, we hypothesized that the lower values from PSA assay reflected incomplete digestion of the dendrimer core. Recent work quantifying the EPS of another fungal species, *Tremella fuciformis*, found that the accuracy of the PSA assay could be dramatically increased by a 30-minute incubation in HCl (18). When we adjusted the PSA assay to include this HCl incubation the two *C. neoformans* strains 24067 and Mu-1 resulted in 1.2 ± 0.2 and 4.0 ± 4.1 mg/mL respectively while the *C. gattii* strains 409 and KT24066 resulted in 2.8 ± 1.8 and 1.5 ± 0.3 mg/mL respectively.

Unfortunately, the PSA assay yields still deviated from the dry weight after a 30-minute HCl incubation. Consequently, we performed a time course of HCl incubation to determine the time needed for maximum yield, but before monosaccharides begin breaking down. For this HCl time course we utilized a matched serotype A set of strains, Mu-1 and KN99 to represent both single-motif and mixed GXM motif expression. We have previously hypothesized that the shed EPS may differ in complexity from the CPS attached to the cell (28, 29), so we included both isolated EPS and CPS prepared as detailed previously (16). The PSA assay yield after different times of HCl incubation varied by strain (Supp. Figure 2). Mixed GXM motif strain KN99 EPS and CPS showed similar PS amounts with a maximum yield after 70 minutes 2 M HCl pre-incubation (EPS = 2.1 mg/mL, CPS = 1.8 mg/mL). Unlike KN99, the single-motif expressing strain Mu-1 EPS and CPS showed different HCl incubation times. The maximum Mu-1 EPS yield was observed after 30 minutes 2 M HCl preincubation at 1.03 mg/mL while Mu-1 CPS yield maximum was observed after 50 minutes preincubation at 1.9 mg/mL.

### GXM ELISA measurements

ELISA with IgM 2D10 to capture and IgG1 18B7 to detect was used to measure GXM content. When the resulting optical densities were compared against an H99 strain EPS standard of known concentration, this method yielded PS concentrations for three of the four single motif expressing strains – 24067 at 3.9 ± 4.4, Mu-1 at 4.3 ± 2.4 and 409 at 13.5 ± 6.1 mg/mL. However, we were unable obtain consistent signal for strain KT24066 which expresses the M4 motif of GXM. Recent work examining antibody affinity by glycan array indicated that another mAb to GXM, IgM 12A1, may prove effective in binding the M4 motif of GXM (20). Using mAb 12A1 we observed signal for KT24066 EPS at 0.9 ± 0.5 mg/mL. Statistical analysis shows a significant difference between the GXM ELISA quantification for *C. gattii* strains 409 and kt24066 EPS (p=0.0391).

### Cryptococcal Clinical Tests show variability in detecting cryptococcal PS

We compared two tests utilized in the clinic, the lateral flow assay (LFA) (IMMY) and the cryptococcal latex agglutination assay (CLAA) (Remel). We utilized these kits-to measure cryptococcal PS following the package instructions but with single motif expressing strain PS solutions in place of patient cerebral spinal fluid. These kits differ by method of testing – strip visualization versus agglutination – as well as the antibodies used. The limit of detection for the LFA varied by strain but showed no statistically significant differences (Figure 3A). *C. neoformans* strains 24067 (0.127 ± 0.002 µg/mL) and Mu-1 (0.314 ± 0.011 µg/mL) had a lower limit of detection than *C. gattii* strains 409 (1.81 ± 3.5 µg/mL) and KT24066 (4 ± 0 µg/mL), with the mixed motif serotype A KN99α strain limit of detection between that of *C. neoformans* and *C. gattii* single motif expressing strains (Figure 3A). On the other hand, the limit of detection of the CLAA was statistically significantly different between each strain (Figure 3B). While strains 24067 (0.667 ± 0.083 µg/mL), Mu-1 (0.833 ± 0.083 µg/mL), 409 (0.667 ± 0.083 µg/mL), and mixed motif strain KN99α all have relatively low limits of detection, kt24066 (10 ± 0 µg/mL) is nearly 10 times greater (Figure 3B).

**Figure 3:**
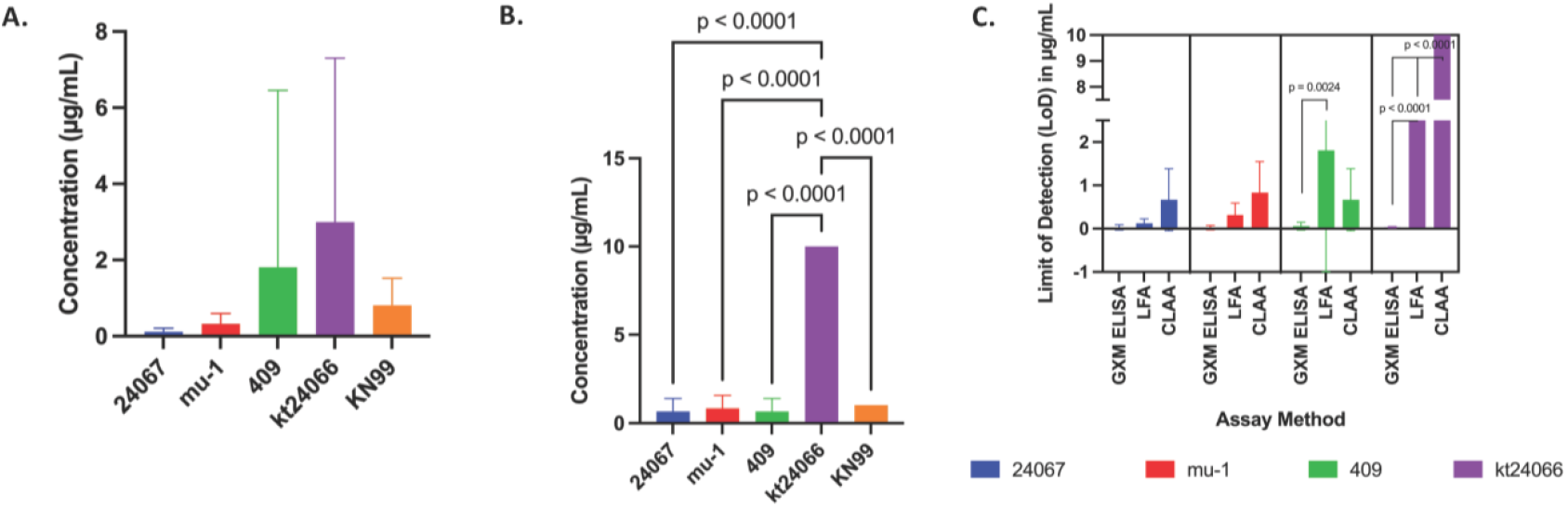
Analysis of Antibody-based Diagnostic tests for Cryptococcal infection using purified EPS. **A**. Lateral Flow Assay **B**. Latex Agglutination Assay **C**. Comparison of three antibody assays for Cryptococcal EPS GXM sandwich ELISA, Latex Agglutination, and Lateral Flow Assay. Statistical analysis of difference between assays, two-way ANOVA with Bonferroni multiple analysis correction.

### Assessment of differences between methods

A goal of this work was to assess the replicability and sensitivity of laboratory and clinical assays for measuring cryptococcal PS. To assess replicability, we examined the variability within each method (Figures 3C and 4). This analysis shows statistically significant variability within two methods, GXM ELISA and clinical assay CLAA (Figure 4B). To assess the sensitivity as a function of GXM motif expression we looked at the variability of method quantification by strain (Figures 3C and 4). Since the laboratory methods yield amount while the clinical assays yield a limit of detection these were assessed separately but with the GXM ELISA included in both. Comparison of GXM ELISA, LFA, and CLAA limit of detection showed no statistically significant difference between methods for *C. neoformans* strains 24067 and Mu-1 (Figure 3C). However, both *C. gattii* strains 409 and kt24066 showed statistically significant differences between methods with only kt24066 showing significant differences between all three methods (Figure 3C). Comparison between strains also showed statistically significant differences between laboratory methods, though no species trend was observed the least variation is seen for Mu-1 (Figure 4). Statistical analysis shows a significant difference between all strains with regards to dry weight and PSA quantification, which has the lowest yields while 409 and kt24066 show significant differences between each method of quantification (Figure 4). The sensitivity of these methods is challenging to compare as most laboratory methods are expressed as a quantification in mg/mL while the clinical assays and GXM ELISA are expressed as limits of detection in µg/mL.

**Figure 4:**
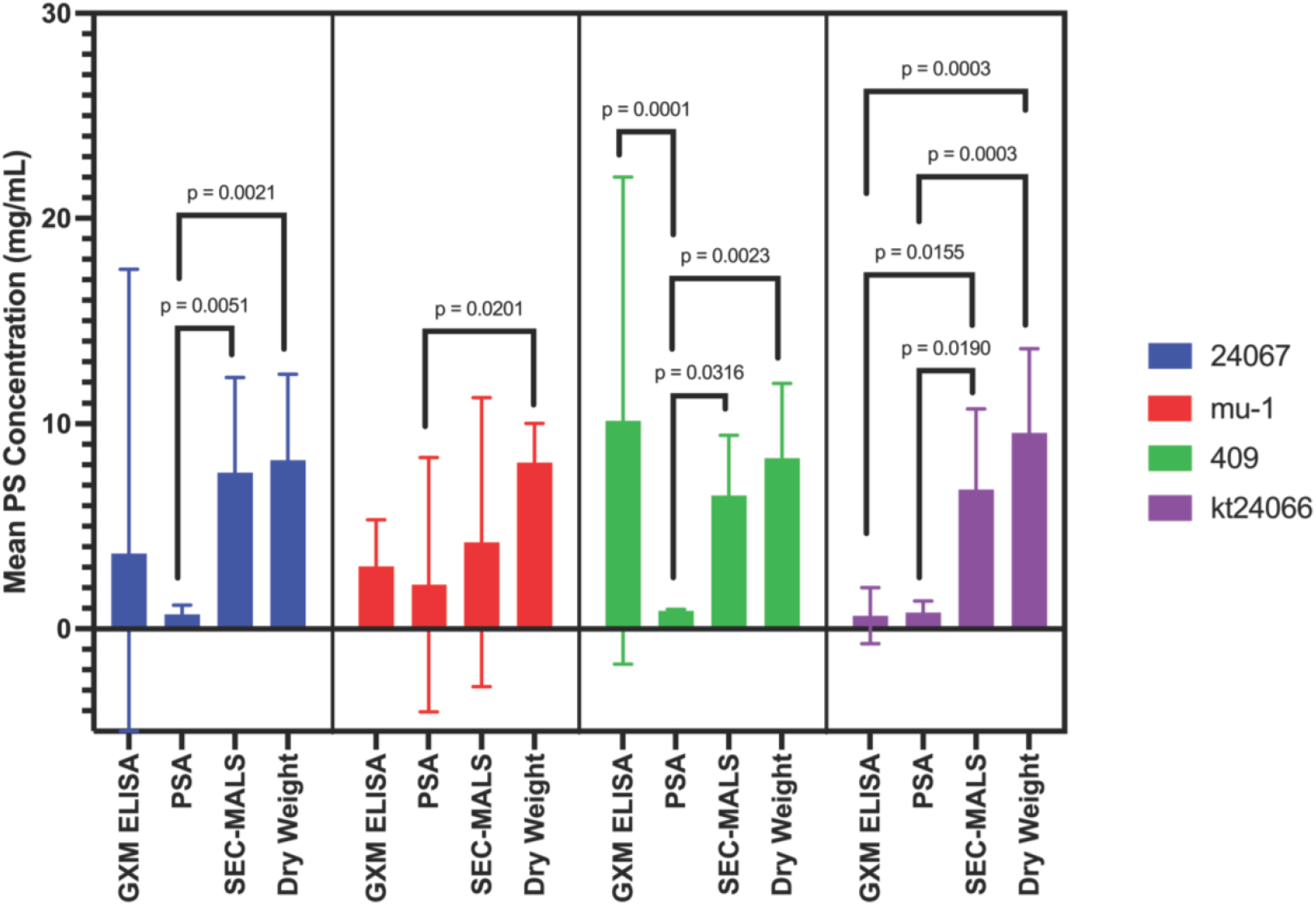
Comparison of four polysaccharide quantification methods. For each strain, statistical analysis comparison between quantification method using two-way ANOVA with Bonferroni correction for multiple comparisons.

## Discussion

The goal of this study was to assess the methods of polysaccharide quantification with regards to, replicability, GXM motif specificity, and sensitivity. These comparisons show that only two methods have reduced replicability, GXM ELISA and clinical assay CLAA. The variability of quantification by strain allows us to assess the GXM motif detectability of these methods. The specificity of all methods was best for strain Mu-1 which expresses the M2 GXM motif and worst for strain kt24066 which expresses the M4 GXM motif. Interestingly, the antibody-based assays GXM ELISA, LFA and CLAA also show a sensitivity difference by cryptococcal strain. All three methods show similarly high sensitivity for *C. neoformans* strains 26067 and Mu-1 expressing GXM motifs M1 and M2 respectively. However, the GXM motif specificity of GXM ELISA is higher for *C. gattii* strains 409 and kt24066, expressing GXM motifs M3 and M4 respectively, than the LFA and CLAA clinical assays. These antibody-based assays, while showing variability in their sensitivity for single GXM motif PS, are the most sensitive with the GXM ELISA allowing for sample measurement of PS <1 µg/mL.

These comparisons show that dry weight had the best replicability while the GXM ELISA was most sensitive. Overall, all assays showed the highest sensitivity for the M2 GXM motif, though this could be affected by affinity of the antibodies used in the antibody-based methods. Additionally, we would note that SEC-MALS shows high replicability and yields the most information about PS. While the sensitivity of the PS quantification method varied by desired attributes, we found other observations about these assays to be pertinent and relevant to the cryptococcal field at large.

The PSA assay yielded a surprising result with regards to EPS and CPS. We observed that the HCl pre-incubation times for the PSA assay varied while both strains are *C. neoformans* and serotype A, the KN99 strain is known to express multiple GXM motifs while the Mu-1 strain only expresses only the M2 motif. Therefore, we were surprised to note that while Mu-1 EPS required 30-minute HCl incubation, as reported for other polysaccharides in the literature (18), both Mu-1 CPS and both KN99 EPS and CPS required longer HCl pre-incubation to reach maximum yield. Additionally, while Mu-1 CPS max yield was observed after 50 minutes HCl pre-incubation, the maximum yield for both EPS and CPS from KN99 took 70 minutes HCl pre-incubation. The HCl pre-incubation should chemically cleave bonds linking sugar residues within the polysaccharide that contribute to higher order structure, suggesting that the more complex the structure of the polymers, the longer the required HCL pre-incubation. This strongly suggests that EPS and CPS, at least in single motif expressing strains have differing levels of structural complexity. Additionally, it suggests that the expression of multiple GXM motifs results in more complex higher order PS structures than single GXM motif expression. This adds to accumulating evidence suggesting cryptococcal EPS and CPS are distinct species (28, 30). The observation that the more structurally complex PS required longer HCL digestion suggests that as complexity increases it takes longer for hydronium ions to find sugar linkages for hydrolysis.

We noted issues with some of the assays, specifically related to antibody use. First was the antibody set used for ELISA (13) are mAbs IgG1 18B7, and IgM 2D10. These antibodies were raised against NIH-371 EPS tetanus toxoid (TT) conjugate (31), while 18B7 binds the M1, M2, and M4 motifs 2D10 binds only the M2 motif (20), despite previous reports that 2D10 binds M1 motif EPS in ELISA (13, 32). While the dry weight analysis yielded an average of 9.5 mg/mL, multiple replicate ELISAs showed no signal for kt24066 EPS. After switching the IgM antibody to 12A1 we were able to obtain an average yield of 0.64 mg/mL by ELISA for kt24066 EPS, but this is still far less than the dry weight yield or even the SEC-MALS yield of 6.79 mg/mL. This is clearly due to kt24066 expressing only the M4 motif of GXM, which is not found in the EPS of strain NIH-371 which was used to make the GXM-TT from which all three antibodies originated. To determine if this was just our set of anti-GXM mAbs or more widespread, we utilized the ELISA-derived PS concentrations to determine the limit of detection for each strain by both LFA and CLAA. The limit of detection for 409 (M3) and kt24066 (M4) EPS is higher in both the LFA and CLAA, though only statistically significantly so for kt24066 in the CLAA.

In performing both the LFA and CLAA clinical assays we noted prozone effects. For the CLAA clinical assay we observed a binding range from 0.5-10 µg/mL for 24067 (M1), Mu-1 (M2) and 409 (M3) and a range of 10 µg/mL and greater for kt24066. The CLAA clinical assay utilizes the mAb IgM BA4-DF5 which resulted from immunization with, and was shown to bind, strains not expressing the M3 or M4 motifs of GXM. In contrast the LFA clinical assay shows much broader binding ranges for all four single motif expressing strains. 24067 (M1) has a range of 0.13-78 µg/mL, Mu-1 (M2) 0.3-385 µg/mL, 409 (M3) 1.8-252 µg/mL and KT24066 (M4) 3-640 µg/mL. These ranges are not surprising given that the LFA clinical assay utilizes two mAbs F12D2 which resulted from immunization with the Mu-1 (M2) strain and 339 which resulted from immunization with 24065 (M3). Both antibodies were shown to bind strains expressing motifs 1, 2, 3, 5, and 6. Clearly the antibodies from both the CLAA and LFA clinical assays bound the M4 motif of GXM, but not with the affinity which they bind motifs M1-M3.

The antibodies utilized for ELISA, CLAA, and LFA all resulted from immunization with different cryptococcal strains, though all contained motifs M1, M2, M5, and M6. These antibodies have a lower affinity for the M4 motif which has structural similarities to both the M3 and the M5 motifs (33). While we cannot confirm the affinity range for the M5 GXM motif, the reported binding to M5 and clear lack of binding to M4 suggests the nearly fully doubly substituted nature of the M4 motif induces steric hinderance inhibiting the binding of most anti-GXM mAbs. Future work will be necessary to determine the exact nature of M4 binding inhibition. Given the dependance of anti-GXM mAb binding on GXM Man-C6-o-acetylation (21, 34, 35), the acetylation patter of GXM motifs will also need to be determined to rule this out of the mechanism of M4 binding inhibition.

While many clinical isolates of *Cryptococcus* spp. undergo molecular characterizations (36–40), they rarely include GXM motif characterization. Consequently, information about this critical virulence factor is usually unavailable when diagnosing and treating cryptococcal infections. Historically capsule characterizations have relied on serotype in cryptococcal disease diagnostics, within serotype A, that which is causative in the majority of cryptococcal infections, strains can express a single GXM motif, or all of them (14). While the proportion of single motif strains among clinical specimens is unknown these represent a minority (14%) of the strains for which GXM has been structurally characterized. Of the thousands of cryptococcal strains for which molecular type or genotype are available, GXM motif expression has been described for less than one hundred. The work detailed here shows substantial variability in anti-GXM antibody binding to different GXM motifs. Across the antibody-based major clinically diagnostic tests, binding to two serotype A strains were dramatically different. The LFA showed a range of 1,000-0.811 µg/mL for KN99 while the CLAA showed a range of 10-1 µg/mL. For Mu-1 the signal range for the LFA was 385-0.31 µg/mL and 10-0.5 µg/mL for the CLAA. Additionally, for the LFA assay while the prozone effect was noted for Mu-1 over the concentration of 385 µg/mL, no prozone effect was observed for KN99 even at 1 mg/mL. This range of binding within just serotype A was unexpected, though we must note that the range in binding across the four serotypes and GXM motifs examined here is significantly larger than within serotype A.

Our observations that LFA and CLAA produce different results with strains expressing different GXM repeat motifs could have significant clinical implications. Certainly, equivalent LFA and CLAA titers for two patients infected with different cryptococcal strains do not necessarily imply that each has the same PS content in their serum or cerebrospinal fluid. Given that PS has profound deleterious effects on the immune response the lack of comparability for titers as an indicator of GXM motif or amount of PS could confound attempts to generalize aspects of cryptococcal pathogenesis and their applicability to individual patients. At the very least, our observations warrant a more detailed exploration of the sensitivity and specificity of these clinical tests for cryptococcal strains expressing structurally variable polysaccharides to understand the potential for false negative results because of strain antigenic variation.

In summary, quantitation of cryptococcal PS is method dependent. For laboratory experiments dry weight and SEC-MALS are probably the most reliable methods with the caveat that neither is as sensitive as the serological assays used clinically. However, the serological assays revealed significant strain-to-strain variation, a finding consistent with fact that the PS from different strains can differ in epitope content/accessibility and structural complexity. Our findings indicate a complex relationship between polysaccharide structure and the reliability of the various methods used for its quantitation and suggest the need for further investigations into these problems.

## Acknowledgements

This work was supported by R01AI152078 and R01AI162381 from the National Institutes of Allergy and Infectious Diseases awarded to A.C.

## Footnote page

## Any meeting(s) where the information has previously been presented

This work has not been presented at any scientific meetings.

**Table 1:** Summary of data collected for PS Quantification

| **Method** | **Strain** | **Replicate** |  |  | **Analysis** |  |
| --- | --- | --- | --- | --- | --- | --- |
| **ELISA**<br>**(mg/mL)** | **Strain** | **Rep 0** | **Rep 1** | **Rep 2** | **Average** | **Std. Dev.** |
|  | 24067 | 0.78 | 0.14 | 10.09 | 3.67 | 4.55 |
|  | Mu-1 | 3.85 | 2.05 | 3.24 | 3.05 | 0.75 |
|  | 409 | 10.1 | 5.39 | 14.94 | 10.14 | 3.90 |
|  | Kt24066* | 0.64 | 0.09 | 1.19 | 0.64 | 0.45 |
| **PSA**<br>**(mg/mL)** | **Strain** | **Rep 0** | **Rep 1** | **Rep 2** | **Average** | **Std. dev** |
|  | 24067 | 0.72 | 0.87 | 0.51 | 0.70 | 0.15 |
|  | Mu-1 | 0.56 | 0.86 | 5.03 | 2.15 | 2.04 |
|  | 409 | 0.89 | 0.90 | 0.84 | 0.88 | 0.03 |
|  | kt24066 | 0.63 | 1.05 | 0.68 | 0.78 | 0.19 |
| **SEC-MALS**<br>**(mg/mL)** | **Strain** | **Rep 0** | **Rep 1** | **Rep 2** | **Average** | **Std. dev** |
|  | 24067 | 9.30 | 5.63 | 7.93 | 7.62 | 1.51 |
|  | Mu-1 | 1.05 | 5.11 | 6.52 | 4.23 | 2.32 |
|  | 409 | 5.82 | 5.84 | 7.87 | 6.51 | 0.96 |
|  | kt24066 | 7.79 | 4.96 | 7.62 | 6.79 | 1.29 |
| **MW (kDa)** | **Strain** | **Rep 0** | **Rep 1** | **Rep 2** | **Average** | **Std. dev** |
|  | 24067 | 8.81 | 179.80 | 99.31 | 95.97 | 69.85 |
|  | Mu-1 | 142.30 | 259.40 | 120.90 | 174.20 | 60.87 |
|  | 409 | 7.22 | 190.50 | 90.13 | 95.95 | 74.94 |
|  | kt24066 | 94.68 | 283.70 | 41.89 | 140.09 | 118.75 |
| **Dry Weight**<br>**(mg/mL)** | **Strain** | **Rep 0** | **Rep 1** | **Rep 2** | **Average** | **Std. dev** |
|  | 24067 | 8.00 | 6.67 | 10.00 | 8.22 | 1.37 |
|  | Mu-1 | 7.67 | 7.67 | 9.00 | 8.11 | 0.62 |
|  | 409 | 7.33 | 7.67 | 10.00 | 8.33 | 1.19 |
|  | kt24066 | 10.33 | 10.67 | 7.67 | 9.56 | 1.34 |

**Table 2:** Summary of data collected for antibody assays, limit of detection

| Method | Strain | Replicate |  |  | Analysis |  |
| --- | --- | --- | --- | --- | --- | --- |
| LFA (µg/mL) |  | R0 | R2 | R3 | Average | Std. Dev. |
|  | **24067** | 0.15 | 0.15 | 0.08 | 0.13 | 0.04 |
|  | **Mu-1** | 0.19 | 0.38 | 0.38 | 0.31 | 0.09 |
|  | **409** | 0.49 | 0.99 | 3.95 | 1.81 | 1.53 |
|  | **kt24066** | 4.00 | 4.00 | 4.00 | 4.00 | 0.00 |
| Agglutination (µg/mL) |  | R0 | R2 | R3 | Average | Std. Dev. |
|  | **24067** | 0.5 | 1 | 0.5 | 0.67 | 0.24 |
|  | **Mu-1** | 0.5 | 1 | 1 | 0.83 | 0.24 |
|  | **409** | 0.5 | 1 | 0.5 | 0.67 | 0.24 |
|  | **kt24066** | 10 | 10 | 10 | 10.00 | 0.00 |
| GXM ELISA (µg/mL) |  | R0 | R2 | R3 | Average | Std. Dev. |
|  | **24067** | 0.03 | 0.003 | 0.05 | 0.03 | 0.02 |
|  | **Mu-1** | 0.04 | 0.002 | 0.03 | 0.02 | 0.02 |
|  | **409** | 0.09 | 0.02 | 0.07 | 0.06 | 0.03 |
|  | **kt24066** | 0.04 | 0.03 | 0.03 | 0.03 | 0.00 |

