## Supplemental Figures for "Unexpected variability in laboratory and clinical assays used to quantify *Cryptococcus neoformans* polysaccharide"

Supplemental Data

Supplemental Figure 1

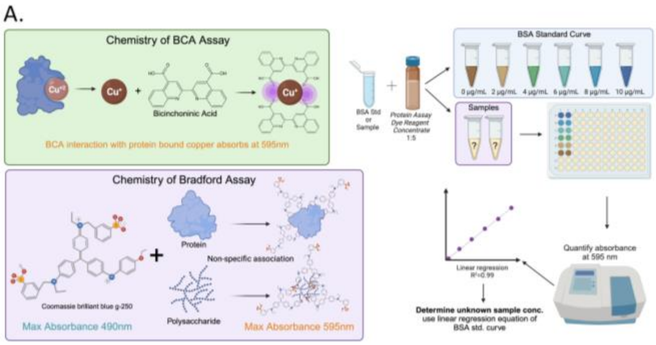

B.

| | | 24067 EPS | Mu-1 EPS | 409 EPS | kt24066 EPS |
|---|---|---|---|---|---|
| Rep #0 | Protein µg/mL | 6.78 | 7.12 | 8.08 | 8.38 |
| | PS mass µg/mL | 8000.00 | 7666.67 | 7333.33 | 10333.33 |
| | % protein | 0.08% | 0.09% | 0.11% | 0.08% |
| Rep #1 | Protein µg/mL | 8.32 | 9.10 | 9.30 | 8.77 |
| | PS mass µg/mL | 6666.67 | 7666.67 | 7666.67 | 10666.67 |
| | % protein | 0.12% | 0.12% | 0.12% | 0.08% |
| Rep #2 | Protein µg/mL | 5.44 | 5.69 | 6.69 | 6.37 |
| | PS mass µg/mL | 10000.00 | 9000.00 | 10000.00 | 7666.67 |
| | % protein | 0.05% | 0.06% | 0.07% | 0.08% |
| | **Avg % protein** | **0.09%** | **0.09%** | **0.10%** | **0.08%** |

**Supplemental Figure 1:** Quantification of EPS-associated protein by BCA assay. **A.** Overview of the chemistry of two predominant protein quantification assays, BCA and Bradford (left) showing non-specific Coomassie interaction allows for interaction with polysaccharides. Method of BCA protein quantification (right). **B.** Protein quantification in µg/mL as well as representative percentage of total sample mass for three biological replicates of four SMES EPS.

Supplemental Figure 2

A.

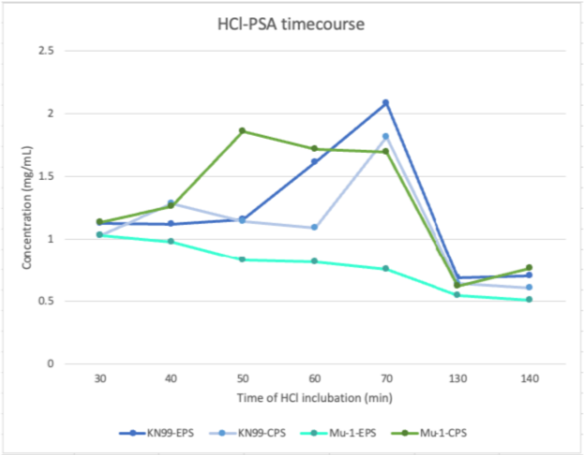

| Strain | PS Type | 30 min | 40 min | 50 min | 60 min | 70 min | 130 min | 140 min |
|---|---|---|---|---|---|---|---|---|
| KN99 | EPS | 1.129 | 1.119 | 1.159 | 1.611 | **2.086** | 0.689 | 0.705 |
| | CPS | 1.028 | 1.284 | 1.142 | 1.090 | **1.811** | 0.548 | 0.511 |
| Mu-1 | EPS | **1.029** | 0.976 | 0.831 | 0.813 | 0.752 | 0.639 | 0.605 |
| | CPS | 1.137 | 1.261 | **1.858** | 1.717 | 1.694 | 0.620 | 0.760 |

B.

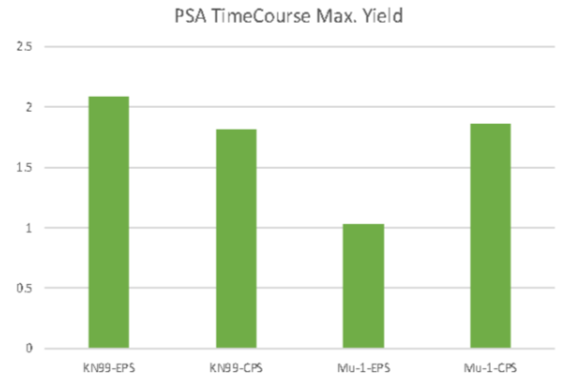

**Supplemental Figure 2:** PSA quantification after HCl incubation for cryptococcal polysaccharide quantification. **A.** Graph of PSA quantification after incubation in 4N hydrochloric acid. **B.** PSA quantification of Eps and CPS for single motif expressing strain Mu-1 (M2) and mixed motif expressing strain KN99 throughout time course. Bolded values represent highest PSA value timepoint.
